# Fast purification of recombinant monomeric amyloid-β from *E. coli* and amyloid-β-mCherry aggregates from mammalian cells

**DOI:** 10.1101/2020.05.13.093534

**Authors:** Amberley D. Stephens, Meng Lu, Gabriele S. Kaminski Schierle

## Abstract

The Alzheimer’s disease related peptide, Amyloid-beta (Aβ)1-40 and 1-42, has proven difficult to be purified as a recombinant monomeric protein due its expression in *E. coli* leading to the formation of insoluble inclusion bodies and its tendency to quickly form insoluble aggregates. A vast array of methods have been used so far, yet many have pitfalls, such as the use of tags for ease of Aβ isolation, the formation of Aβ multimers within the time frame of extraction or the need to reconstitute Aβ from a freeze dried state. Here, we present a rapid protocol to produce highly pure and monomeric recombinant Aβ using a one-step ion exchange purification method and to label the peptide using a maleimide dye. The solublisation and purification steps take only three hours. We also present a protocol for the isolation of Aβ-mCherry from mammalian cells.

**Highlights:**

- Purification of untagged, monomeric recombinant Aβ from *E. coli*.
- A fast protocol; 6 hours for *E. coli* growth and Aβ expression, 2 hours to clean inclusion bodies, 45 mins to solublise and purify the peptide.
- No freeze drying step that can lead to oligomer formation.
- Purification of fluorescent Aβ-mCherry from mammalian cells.

## Introduction

The presence of Amyloid-beta (Aβ) plaques and Tau tangles in neurons are hallmarks of Alzheimer’s disease, therefore great research effort is put towards understanding how these initially soluble proteins misfold and contribute to pathology. To study protein misfolding, large quantities of protein are required, and while purification protocols for Tau proteins are fairly well established, those for Aβ, in its isoforms of 1-39/43 and its mutant variants, are very heterogeneous and lead to variable products ^1^.

Many studies investigating aggregation rates and toxicity of Aβ currently use synthetic Aβ due to the ease of purchase and the little handling required to obtain monomeric Aβ. However, Aβ can be expensive to purchase and needs to be reconstituted to remove oligomers, leading to loss of protein and variation in the resulting sample due to sample impurities^2^ and due to use of different reconstitution protocols, such as the use of hexafluoroisopropanol (HFIP) or ammonium hydroxide^3^. Furthermore, presence of impurities in synthetic Aβ can influence aggregation propensity and toxicity^4^.

Purification of tagged-Aβ is a highly popular method as addition of tags can improve solubility and permit the use of affinity capture chromatography which can yield highly pure recombinant Aβ samples. In a recent review on Aβ purification methods, it was highlighted that 23/30 protocols utilised tagged-Aβ for purification^1^. The added benefit of using a recombinant tagged-system containing a cleavage site at the Aβ N-terminus is that it can be utilised to release the wild-type Aβ sequence without a methionine (M) start codon. *In vivo*, Aβ is cleaved from the amyloid precursor protein, therefore the first codon in the sequence is an aspartate, the sequence of which cannot be obtained by expressing Aβ alone, and AβM variants are instead used which have the methionine starting residue before aspartate. However, if tags are not removed prior to further analysis of the peptide even a small tag, such as a 6xHis-tag, can greatly influence the protein structure and aggregation propensity^5^. Moreover, removal of the tag requires the addition of a cleavage recognition site and additional purification steps which lead to loss of protein, increased time of protein handling and therefore to the formation of aggregated species.

The protocol by Walsh et al., provided an easy method for purification of AβM variants using urea solubilisation to isolate AβM from inclusion bodies, purification by ion exchange chromatography and size exclusion chromatography, centrifugation applying a 30 kDa filter and lyophilisation to store the recombinant protein^6^. However, the DEAE-cellulose chromatography media used in the ion exchange step is no longer commercially available.

Reversed phase (RP) chromatography is another frequently used method to purify Aβ due to the high purity of the resulting recombinant protein. Aβ is eluted from the RP column along a gradient of organic solvent in the presence of an ioniser such as trifluoracetic acid (TFA). The organic solvent is then removed by freeze drying. The process of freeze drying induces formation of oligomers, which subsequently can be removed by gel filtration, yet this adds another step to the purification protocol. We have shown for another amyloidogenic protein, α-synuclein, that freeze drying leads to a compaction of the monomer structure and formation of heterogeneously sized oligomers, even after reconstitution in buffer, compared to samples that were frozen directly after purification^7^. Freeze drying could also affect the structure of monomeric Aβ, although further studies are needed to confirm this. As with all intrinsically disordered amyloidogenic proteins the structure of the starting material e.g. presence of multimers or degraded products, and the surrounding environment heavily influence the aggregation rate, the pathways of aggregation that are taken, and the toxicity of the resulting amyloid^2^.

Here, we provide a fast protocol for the purification of recombinant AβM42 and AβMC40 from *E*.*coli* using a one-step ion exchange chromatography protocol from cleaned and solubilised inclusion bodies. The protocol can be amended to permit the incorporation of a maleimide dye label to the cysteine residue of mutated sequences, such as the AβMC40 sequence. After induction of AβM expression in *E. coli* the purification protocol takes only around three hours to obtain highly pure monomeric AβM. We also present a protocol for the isolation of AβM(E22G)-mCherry from HEK293 cells again using one-step ion exchange chromatography.

## Results

### Thorough washing of inclusion bodies yields pure recombinant AβM

Aβ42 and Aβ40 are the most abundant Alzheimer’s disease-associated variants and will be used here to demonstrate the purification method^8^. The Aβ40 variant used in this protocol contains an additional N-terminus cysteine residue, AβMC40, which can be dye-labelled with thiol reactive dyes. AβM42 and AβMC40 were expressed from pET3a plasmids in *E. coli* BL21 (DE3) pLysS strain with 1 mM IPTG for 4 hours. The *E. coli* cultures were centrifuged in 50 mL falcon tubes and the pellet stored at -80°C until needed (Figure 1, light blue box). During expression, AβM peptides form into cytoplasmic insoluble inclusion bodies, which can be beneficial for the purification process as inclusion bodies can contain highly pure levels of the protein of interest^9^. The inclusion bodies were thoroughly washed to obtain very pure inclusion bodies (Figure 1, mid blue box). The frozen pellet from 50 mL of *E. coli* culture was resuspended in wash buffer 1 (Table 1) which contained protease inhibitors to reduce proteolytic cleavage of AβM during cell lysis, 1 M guanidine hydrochloride (GuHCl) to increase washing efficiency by aiding the removal of other proteins, and 1% triton X-100 to remove lipids bound to the inclusion bodies. The *E. coli* cells were sonicated 5 × 30 s on ice and were centrifuged at 10,000 x g to remove cell debris. This was repeated three times with different additives in the wash buffer (Table 1). By the end of wash 4 the pellet contained highly pure inclusion bodies which were white in colour, and which were consequently stored at -80°C until solubilisation and purification.

**Table 1.** Wash buffer and additives used to clean Aβ containing inclusion bodies.

| 10 mM Tris 1 mM EGTA pH 9 |  |  |  |
| --- | --- | --- | --- |
| Wash 1 | Wash 2 | Wash 3 | Wash 4 |
| protease inhibitors | protease inhibitors |  |  |
| 1 M GuHCl | 1 M GuHCl |  |  |
| 1% Triton X-100 | 1% Triton X-100 | 1% Triton X-100 | No additives |

**Figure 1.**
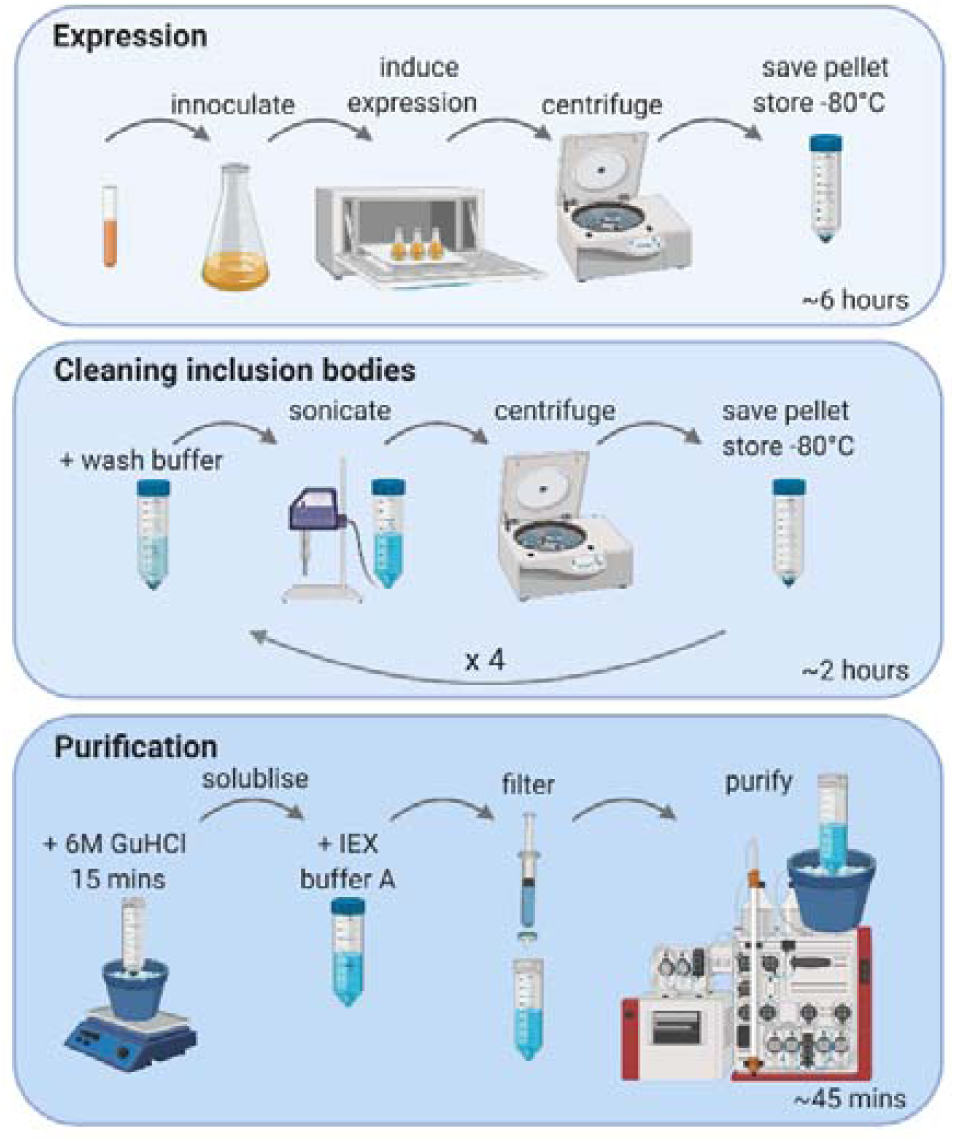
Schematic figure of the expression, isolation and purification of AβM42 and AβMC40 from *E. coli*. *Light blue box, expression* – An overnight culture was inoculated in lysogeny broth (LB) medium and then grown at 37°C until reaching an OD_600_∼0.6-0.8. Expression of AβM was induced upon addition of 1 mM IPTG, four hours after which the culture was centrifuged in 50 mL falcon tubes and the pellet stored at -80°C until use. *Mid blue box, cleaning inclusion bodies* – AβM is retained in insoluble inclusions bodies. The *E. coli* pellet was resuspended in wash buffer 1 (Table 1) and the inclusion bodies were lysed and isolated from the cells by sonication and centrifuged at 10,000 x g. The inclusion bodies were then washed three times with different buffers (Table 1) to remove unwanted proteins and lipids. The final inclusion body pellet was kept at -80°C until use. *Dark blue box, purification* – The pellet of inclusion bodies was solubilised with 200 μL of 6 M GuHCl per 50 mL of culture for 15 minutes on ice on a magnetic stirrer before dilution with 15 mL IEX buffer A. The protein solution was filtered through a 0.22 μm membrane and purified using an ÄKTA FPLC with a HiTrap Q HP ion exchange column.

### One-step ion exchange chromatography yields highly pure monomeric AβM

The AβM was solubilised from the inclusion bodies before purification. 200 μL of 6 M GuHCl was added to the inclusion body pellet from 50 mL of *E. coli* culture on ice. A small stir bar was added and the pellet left on a magnetic stirrer for 15 minutes to solubilise the inclusion bodies. GuHCl was chosen as a solubilising agent instead of urea as urea decomposition leads to isocyanic acid formation which can cause carbamylation of the N-terminus^10^. After 15 minutes, 15 mL of ice cold ion exchange chromatography (IEX) buffer A (10 mM Tris, 1 mM EGTA pH 9) was added to the solution to dilute the GuHCl, reducing the ionic strength of the buffer to permit the protein to bind to the ion exchange column (Figure 1, dark blue box). The protein solution was then filtered through a 0.22 μm filter to remove any precipitate before IEX. The fast protein liquid chromatography (FPLC) machine was not kept in a cold room, therefore all buffers were kept on ice and ice placed around to column to keep the system as cold as possible to reduce AβM aggregation. A HiTrap Q HP column (GE Healthcare) was used to purify the AβM monomer. To keep purification time to a minimum, the column was equilibrated in IEX buffer A prior to sample preparation. AβM was eluted over seven column volumes with a 0-100 % gradient against IEX buffer B (10 mM Tris, 1 mM EGTA, 0.75 M NaCl, pH 9) followed by two column volumes at 100 % buffer B. Absorption at 280 nm was used to monitor protein elution from the column (Figure 2a). Analysis of the eluted fractions by SDS-PAGE on a Coomassie blue stained gel showed pure monomeric AβM eluted at ∼ 30% IEX buffer B (Figure 2, Supplementary Figure 1). The concentration of AβM42 from each 1 mL fraction ranged from 8 - 12.75 μM, determined by absorption at 280 nm and calculated using the extinction coefficient 1490 M^−1^ cm^−1^ (Supplementary Table 1). Purification of the AβMC40 variant is shown in Supplementary Figure 2.

**Figure 2.**
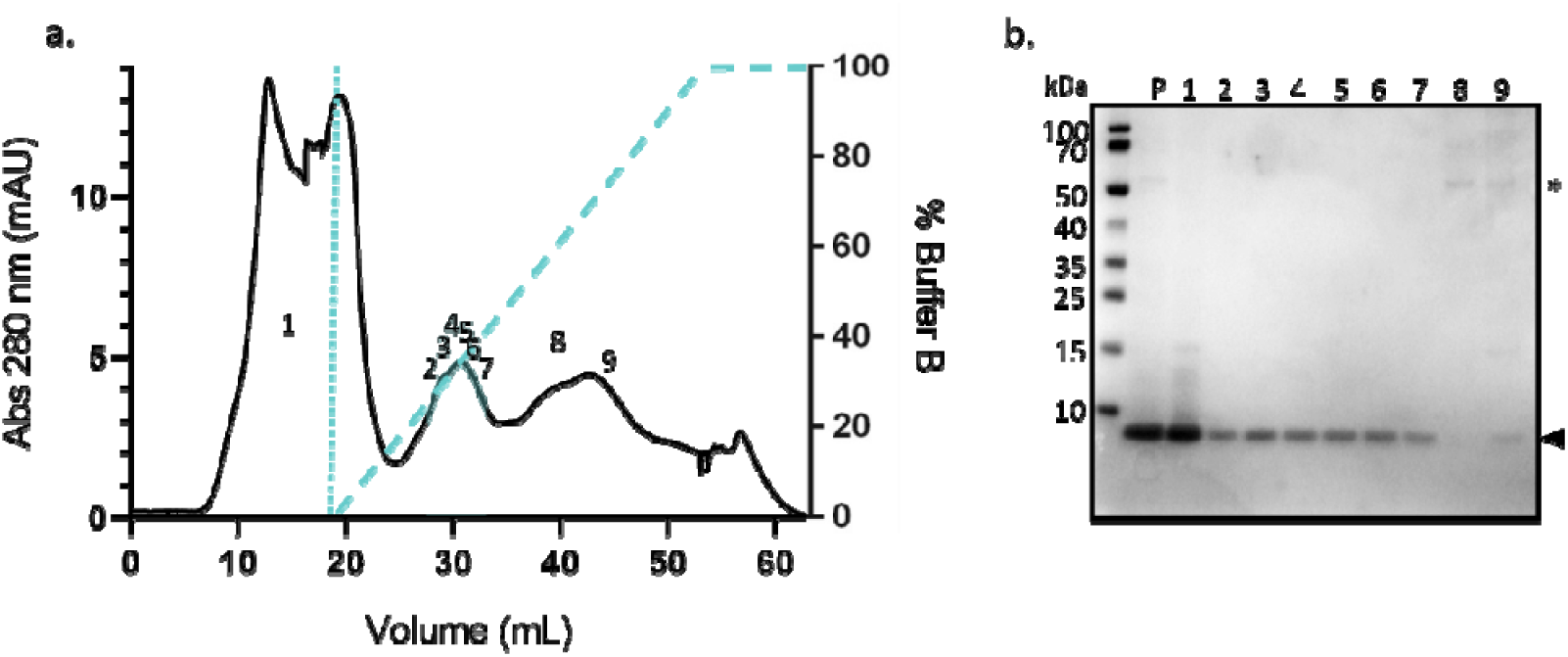
Highly pure monomeric AβM42 is purified by ion exchange chromatography from GuHCl solubilised inclusion bodies. AβM42 was solubilised in 6 M GuHCl and diluted in IEX buffer A before being applied to the HiTrap Q HP ion exchange column (shown up to the first dotted line in a). (a.) The chromatograph of the absorption at 280 nm shows the elution of protein from the HiTrap Q HP column over a gradient of 0-100% buffer B containing 0.75 M NaCl over seven column volumes, followed by two column volumes of 100% buffer B (dashed line showing gradient in a.). (b.) In order to determine when AβM42 got eluted from the column the fractions were collected and analysed using SDS-PAGE on a 4-12% bis-tris gel and Coomassie blue staining. The numbers on the chromatograph a. correspond to the lane on the gel in b. The AβM42 sample prior to IEX (P) was highly pure. Protein bands correlating to ∼ 4.5 kDa (shown by the arrow next to b.) show monomeric AβM42 in fractions 2-7 which are highlighted in blue in the chromatograph (a.). AβM42 eluted at ∼ 30% buffer B. Higher molecular weight species (indicated by a star in b.) elute later in the buffer B gradient in fractions 8 and 9.

### Recombinant AβM42 forms long fibril-like structures over time

The recombinant AβM42 was analysed by liquid chromatography mass spectrometry (LC-MS) to ensure a pure protein of the correct mass/charge had been purified. A deconvoluted mass of 4645 Da was obtained which corresponded to the predicted mass of AβM42 (Figure 3a, m/z data presented in Supplementary Figure 4.). To investigate the aggregation properties of the purified Aβ42, a thioflavin-T (ThT) based aggregation assay was used. The ThT molecule fluoresces when it intercalates into the backbone of a fibril containing β-sheet structure, leading to a sigmoidal curve over time as the protein aggregates and the ThT fluorescence intensity increases11. To investigate aggregation propensity, AβM42 was diluted to 5 μM in 100 mM Tris, 200 mM NaCl, pH 7 with 20 μM ThT and aggregated for 600 mins with a 1 min shake at 300 rpm before every read, every five minutes (Figure 3b). AβM42 aggregation produced a sigmoidal curve of the increase of ThT fluorescence, as expected for a nucleation-dependent protein aggregation assay, with a lag, exponential and plateau phase of increase in aggregation^12^. Shown are AβM42 fibrils formed and imaged by transmission electron microscopy (TEM) (Figure 3c, Supplementary Figure 3).

**Figure 3.**
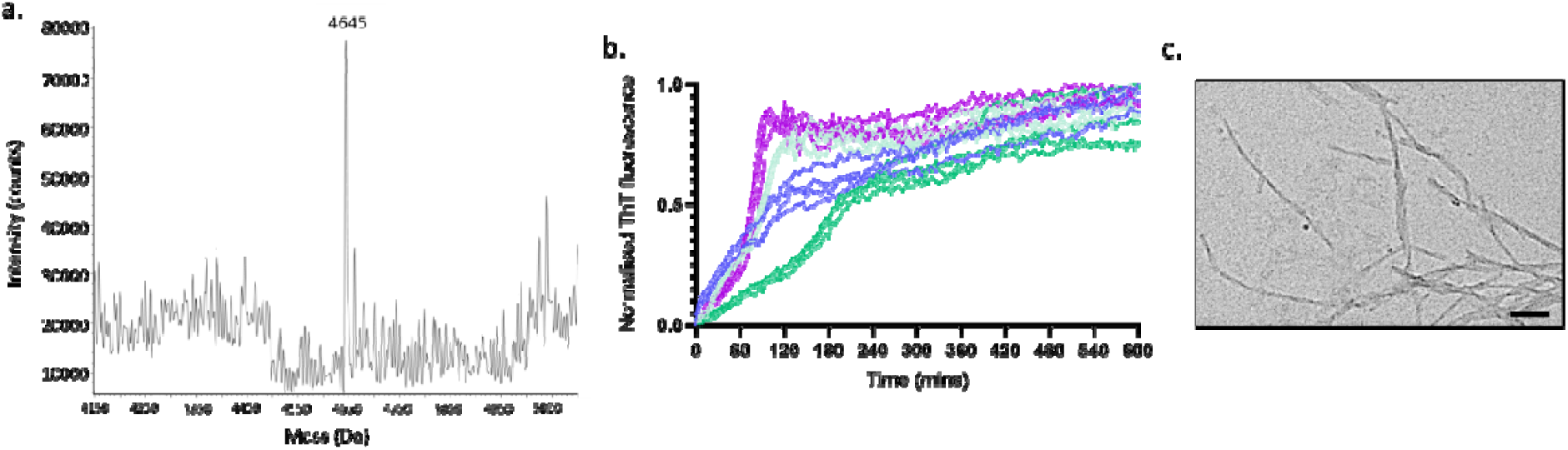
Pure recombinant AβM42 forms long fibrillar structures. Recombinant AβM42 was analysed by mass spectrometry and the (a.) deconvoluted spectrum shows the expected MW of 4645 Da (See Supplementary Figure 4 for m/z spectrum). (b.) Normalized ThT-based aggregation assays show slight batch variation (each colour represents a batch), but good reproducibility within the batches (individual lines represent individual wells, four wells per batch). 5 μM of AβM42 in 100 μM Tris 200 mM NaCl pH 7 with 20 μM of ThT was incubated in a half area 96 well plate at 37°C with double orbital agitation at 300 rpm for one min before each read every five mins for 600 mins. (c) TEM image of fibrils formed during incubation of 5 μM of AβM42 with constant rotation at 20 rpm at 37°C for two days. Scale bar = 100 nm.

### Maleimide dye labelling of AβMC with Alexa Flour 488 leads to fluorescent structures

During the solubilisation step (Figure 1, mid blue box) it is also possible to dye-label a cysteine modified Aβ, where the thiol-reactive dye reacts with the monomeric Aβ as it becomes solubilised. 1 mM tris(2-carboxyethyl)phosphine (TCEP) was added to all buffers to keep the cysteine residue in a reduced form to allow the maleimide dye reaction to occur. All following purification steps remain the same as for the AβM42 purification protocol just described. The AβMC40 monomer labelled with Alexa Flour 488 (AF488), like AβM42, eluted at ∼ 30% of buffer IEX B (Figure 4a). Absorption at 280 nm appears high at ∼60% buffer B (Figure 4a), but as observed on the protein gel only a small amount of high molecular weight species are present (Figure 4b, lanes 7-10, indicated with a star). Instead much of the unbound dye eluted at this point, as observed by the strong dye colour in these fractions and the high absorption at 280 nm was due to interference of the AF488 dye absorption at 280 nm. The concentration of protein and degree of labelling (DOL) was calculated, taking into account the absorption of the dye at 490 nm and the influence of the dye on absorption at 280 nm (Supplementary Table 2). The fluorescently labelled fractions 1, 2, 3 had a concentration of 6.6 μM, 8.9 μM and 8.9 μM, respectively, with a DOL 127.4%, 159.8%, 136.1%, respectively. The DOL may indicate some free dye remaining in the solution as the ratio should be 1:1 for dye to protein. Aggregation of 5 μM of AβMC40-AF488 for two days at 37°C with 20 rpm constant rotation lead to formation of fluorescently labelled fibrils and oligomers (Figure 4c, Supplementary Figure 5).

**Figure 4.**
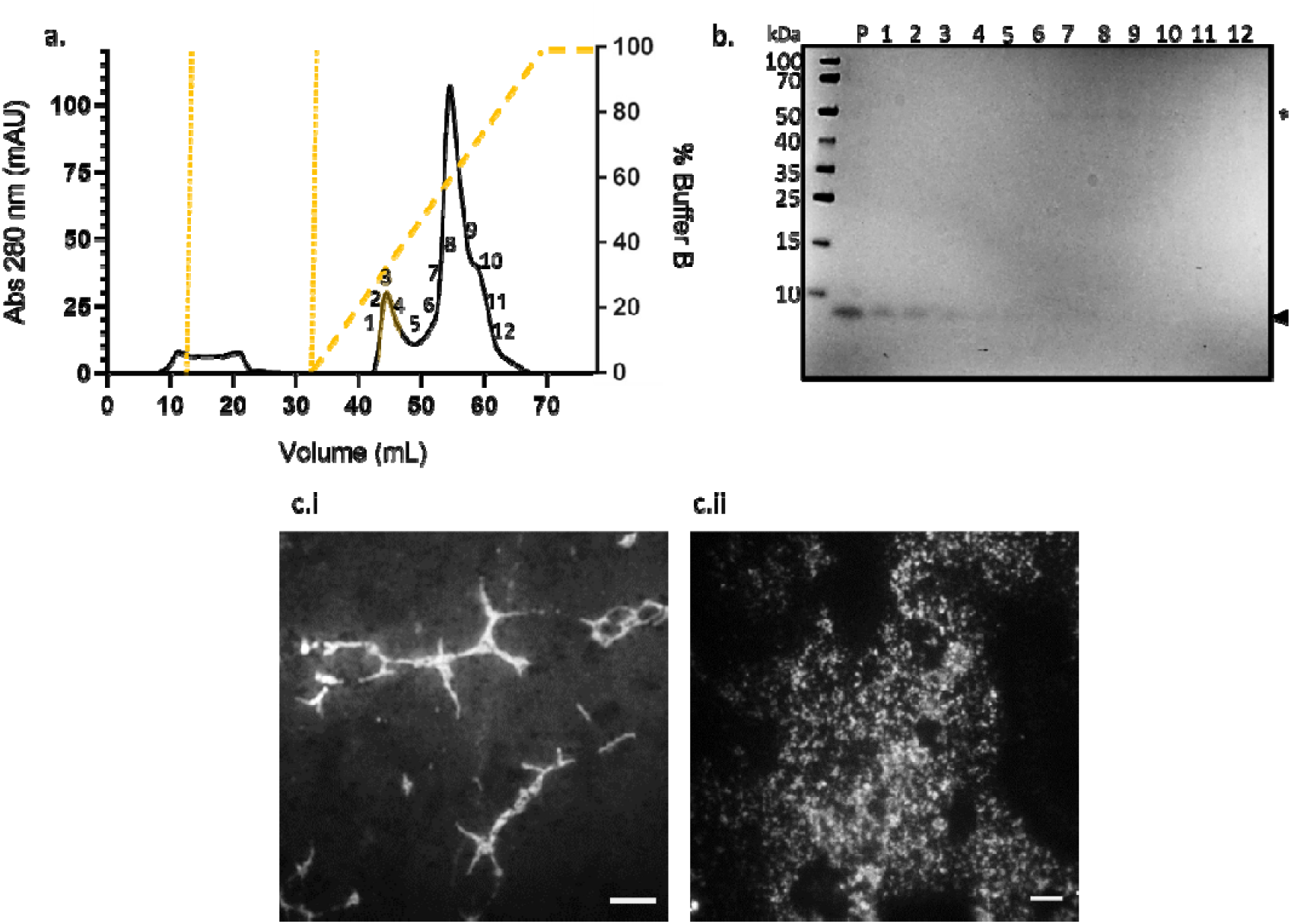
Alexa Flour 488 labelled AβMC40 purified by ion exchange chromatography forms fluorescent fibrils and oligomers. AβMC40-AF488 was solubilised in 6 M GuHCl and diluted in IEX buffer A before being applied to the ion exchange column (shown up to the first dotted line in a.) and unbound protein and dye were washed from the column (shown up to the second dotted line in a.). (a.) The chromatograph of absorption at 280 nm shows the elution of protein from the HiTrap Q HP column over a gradient of 0-100% of buffer B containing 0.75 M NaCl over seven column volumes, followed by two column volumes of 100% buffer B (dashed line showing gradient in a). (b) In order to determine when AβMC40-AF488 got eluted from the column the fractions were collected and analysed using SDS-PAGE on a 4-12% bis-tris gel and Coomassie blue staining. The Aβ sample prior to IEX (P) was highly pure. Protein bands correlating to ∼ 4.5 kDa (shown by the arrow next to b) show monomeric AβMC40-AF488 was present in fractions 1-4, which are highlighted in yellow in the chromatograph a. Higher molecular weight proteins eluting later from the column are indicated by a star in b. (c.) 5 μM of 100% labelled AβMC40-AF488 was incubated for two days at 37°C at 20 rpm and (c.i.) fibrils and (c.ii.) oligomeric/multimeric structures were observed using a widefield microscope. Scale bar 5 μm.

### AβM(E22G)-mCherry purified from HEK293 cells remains fluorescent

While purification of Aβ from *E. coli* is useful as it produces a high yield and quantity, it is still unclear whether the monomeric and/or aggregate structures of Aβ formed *in vitro* are representative of those that form *in vivo*. Here, we present a protocol to isolate AβM(E22G)-mCherry from HEK293 cells. The AβE22G Alzheimer’s disease associated mutant is highly aggregation prone^14^. Although the protein is tagged to a large fluorescent protein, mCherry, which may influence folding and structure, the fluorescent protein permits the study of Aβ aggregation in cells^15^. In this cell line we have previously identified five categories of intracellular AβM(E22G)-mCherry aggregates – oligomers, single fibrils, fibril bundles, clusters and aggresomes. These stages underline the heterogeneity of Aβ42 aggregates and represent the progression of Aβ42 aggregation within the cell ^15^. Isolation of endogenous AβM(E22G)-mCherry may allow further insight into morphology or seeding ability of endogenously structured Aβ by microscopy.

Expression of AβM(E22G)-mCherry was induced by 1 μg/mL tetracycline for seven days. The cells were centrifuged at 4000 x g for 15 mins and the pellet was frozen until use. The cells were lysed by sonication in 50 mM Tris, pH 8 with protease inhibitors before centrifuging to separate the soluble and insoluble fractions. Western blot analysis, probing both Aβ and mCherry, showed that the soluble fraction contained more AβM(E22G)-mCherry than the insoluble fraction (Supplementary Figure 5). The soluble fraction was purified using a HiScreen Capto Q ImpRes ion exchange column and eluted on a linear gradient against IEX buffer B (50 mM Tris, 1 M NaCl pH 8) over seven column volumes, followed by two column volumes of 100% buffer B (Figure 5a). The eluted fractions were analysed by SDS-PAGE separation and the gels were either stained with Coomassie blue (Figure 5b.i) or transferred onto a membrane and probed by Western blot for Aβ (Figure 5b.ii) and mCherry (Figure 5b.iii). The pre-IEX sample (P) contained many proteins (Figure 5b.i), yet a lot appeared to be aggregates of AβM(E22G) (Figure 5b.ii). Unbound proteins (Figure 5a, peak 1, 5b.i, lane 1) eluted from the column during the sample application and column wash. Peak 2, corresponding to lane 2 in Figure 5b. contained the most abundant AβM(E22G) and mCherry by Western blot. The expected MW for monomeric AβM(E22G)-mCherry was 32.3 kDa (Figure 5b.iii, black arrow), therefore both degraded products (Figure 5b.iii, light grey arrow) and aggregated products were present (Figure 5B.ii, star). The presence of Aβ aggregates was to be expected as the E22G mutant is a highly aggregation prone mutant^14^ which was expressed in HEK cells for seven days and no denaturing step was employed during the purification protocol. It appears that mCherry is not always present in Aβ aggregates as the Western blot displaying Aβ bound antibodies is more highly populated than the mCherry Western blot. It possible that the AβM(E22G)-mCherry has become degraded in the cell, the expected molecular weight of mCherry is 26.7 kDa, therefore mCherry fragments may be identified in the Western blot Figure 5b.iii. Another explanation for the discrepancy between the Aβ and mCherry probed blots may be due to steric hinderance preventing the antibody binding to mCherry in an aggregated form. The concentration of fraction 2 was 77 μM, as determined by absorption at 280 nm and calculated using the extinction coefficient of 35870 M^−1^cm ^−1^.

**Figure 5.**
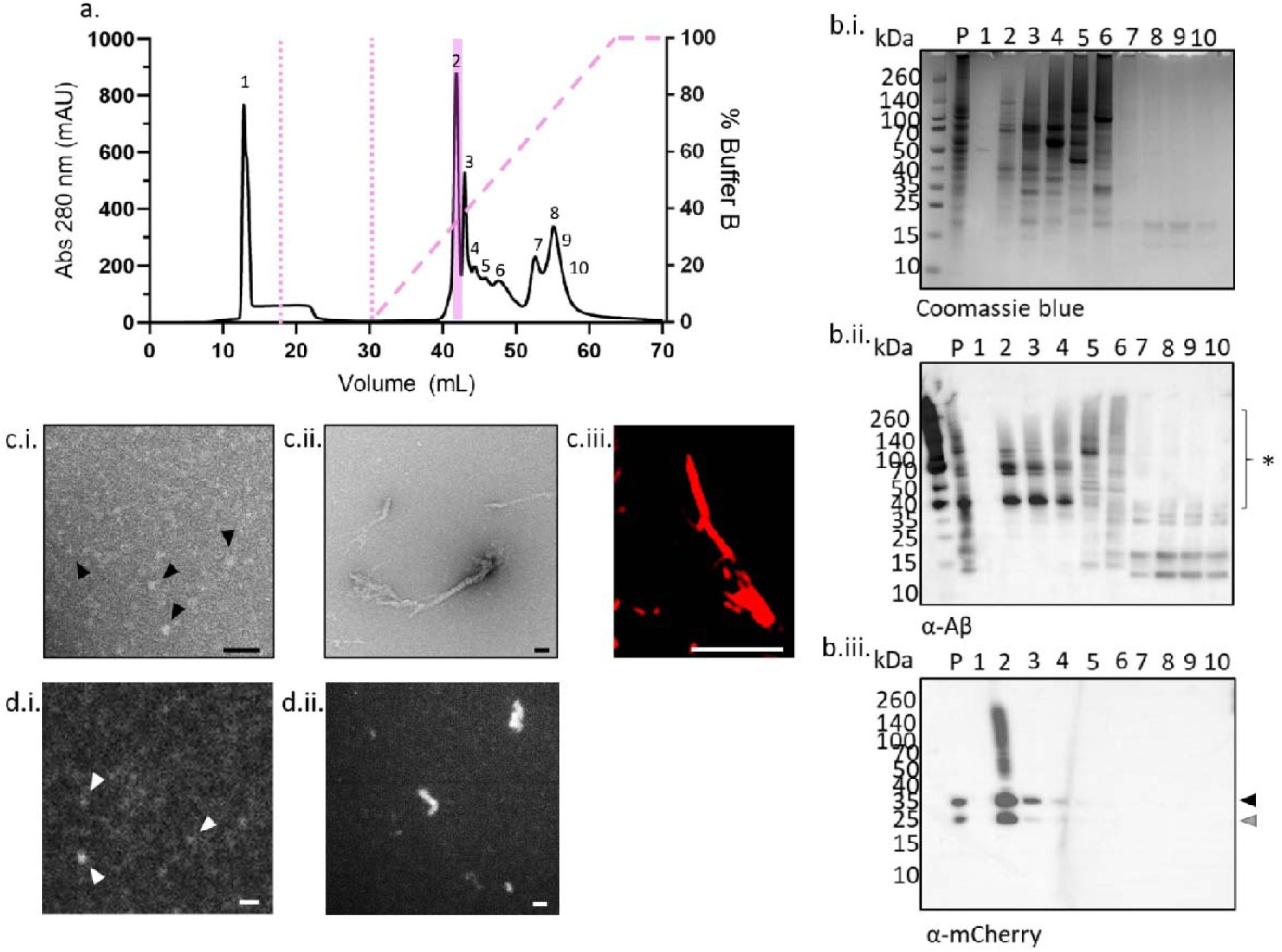
Varying aggregate sizes of AβM(E22G)-mCherry isolated by ion exchange chromatography exhibit weak fluorescence. AβM(E22G)-mCherry lysed from HEK293 cells by sonication. (a.) The soluble fraction was applied to the column (shown up to the first dotted line in a. and the unbound protein was washed from the column (shown up to the second dotted line in a. The chromatograph of absorption at 280 nm show protein elution from the HiScreen Capto Q ImpRes column eluted over a gradient of 0-100% of buffer B containing 1 M NaCl over seven column volumes, followed by two column volumes of 100% buffer B (dashed line showing gradient in a. (b) In order to determine when AβM(E22G)-mCherry eluted off the column, the fractions were collected and analysed using SDS-PAGE on a 4-12% bis-tris gel and (b.i) Coomassie blue staining, or transferred to a membrane for Western blot using antibodies against (b.ii.) Aβ and (b.iii.) mCherry. The sample prior to IEX (P) contained many proteins, including aggregated Aβ and mCherry. Protein bands correlating to ∼ 32.3 kDa (shown by the black arrow in b.iii.) show the predicted MW for monomeric AβM(E22G)-mCherry. Fraction 2 contained the highest content of AβM(E22G)-mCherry, although the presence of degraded mCherry (b.iii., grey arrow) and aggregated Aβ (b.ii., star) were also apparent. The morphology of the purified AβM(E22G)-mCherry was determined by TEM and both (c.i.) oligomers and (c.ii., Supplementary Figure 7) larger aggregates were present. (c.iii.) A section view of an AβM(E22G)-mCherry aggregate inside a cell prior to purification reveals a similar structure to those identified by TEM after purification. These aggregates were also analysed to determine whether they were fluorescent using widefield imaging with a 561 nm laser, both (d.i.) small oligomers and (d.ii.) large aggregates were weakly fluorescent (also see Supplementary Figure 8). Black scale bar = 100 nm, white scale bar = 2 μm.

The morphology and fluorescence of the purified AβM(E22G)-mCherry peak 2 were analysed by TEM and fluorescence microscopy. Both small oligomeric aggregates (Figure 5c.i, indicated by black arrows) and larger aggregates were present (Figure 5c.ii, Supplementary Figure 7.). The large aggregates with fibrillar morphology were very similar to those identified within the cells prior to purification (Figure 5c.iii) ^15^. The aggregates also emitted weak fluorescence when excited with a 561 nm laser (Figure 5d.i., oligomers indicated with white arrows and Figure 5d.ii, shows larger fluorescent aggregates, Supplementary Figure 8.). Mass spectrometry analysis of AβM(E22G)-mCherry showed the expected weight of the monomeric AβM(E22G)-mCherry, 32.3 kDa, was not highly abundant, but that a smaller degraded product of 25.2 kDa and a larger product of 36.3 kDa were the dominant species, amongst many other species of differing molecular weights (Supplementary Figure 9.). In cells Aβ is commonly degraded or altered by post translational modification therefore presence of species of differing molecular weight of AβM(E22G)-mCherry may be reflective of different truncations and modifications that occur within a cellular environment ^13^.

## Conclusion

We present a fast method for the purification of the Alzheimer’s disease related peptide AβM. The protocol can be rapidly completed and if the protein and buffers are kept ice cold, highly pure monomeric recombinant AβM can be obtained. The protocol can be stopped and the products frozen at two key points, after centrifugation of *E. coli* expressing Aβ, or after cleaning of the inclusion bodies prior to solubilisation and purification. Ideally, AβM would be purified and used straight away to prevent freezing of monomeric peptide which can induce dimer and oligomer formation. The purification protocol provided requires only 45 minutes to solubilise and for purification to obtain monomeric AβM. Furthermore, we provide a protocol to isolate fluorescent AβM(E22G)-mCherry structures from mammalian cell lines which can be used to track seeding of Aβ in cells with fluorescent endogenously structured protein.

## Methods and Materials

### Expression of recombinant AβM variants in *E. coli*

The plasmid pET3a containing human AβM42 and AβMC40 cDNA was transformed into *Escherichia coli* (*E. coli*) One Shot^®^ BL21 (DE3) pLysS (Thermo Fisher Scientific, USA). The plasmids were a kind gift from Prof. Sara Linse. The protein sequences encoded in the pET3a plasmids are, AβM42: MDAEFRHDSGYEVHHQKLVFFAEDVGSNKGAIIGLMVGGVVIA and AβMC40: MCDAEFRHDSGYEVHHQKLVFFAEDVGSNKGAIIGLMVGGVV. 1 L cultures of *E. coli* in Lysogeny Broth (LB) containing carbenicillin (100 µg/mL) were grown at 37°C with constant shaking at 250 rpm and induced for expression of AβM when the OD_600_ reached 0.6-0.8 after addition of 1 mM isopropyl-β-thiogalactopyranoside (IPTG). After four hours of AβM expression, the cells were pelleted by centrifugation in 50 mL falcon tubes at 4 k x g for 15 mins. The supernatant is discarded and at this point the pellets can be frozen until further use.

### Cleaning of AβM containing inclusion bodies

30 mL of wash buffer 1 (Table 1) was added to each *E. coli* pellet from 50 mL of culture. 50 mL of culture from a 4 hour induction was found to be a suitable amount that for purification of monomeric AβM, as increasing the concentration can lead to aggregation during purification. At this point, multiple 50 mL pellets can be cleaned at the same time and frozen prior to solubilisation and purification. The pellet was resuspended in 30 mL wash buffer 1 containing 10 mM Tris, 1 mM EDTA, protease inhibitor tablets (cOmplete™ EDTA-free cocktail, Roche), 1 M GuHCl, 1% Triton-X100, pH 9, and was sonicated on ice for 30 s on, 30 s off, five times using an XL-2020 sonicator (Heat Systems, USA). The suspension was centrifuged at 4 °C at 10 k x g for 15 mins. The supernatant was discarded and the pellet was resuspended in 30 mL wash buffer 2 (Table 1) before sonicating it on ice for 30 s on, 30 s off, three times. The suspension was centrifuged at 4 °C at 10 k x g for 15 mins. The supernatant was discarded and the pellet was resuspended in 30 mL wash buffer 3 (Table 1) before being sonicated on ice for 30 s on, 30 s off, three times. The suspension was centrifuged at 4 °C at 10 k x g for 15 mins. The supernatant was discarded and the pellet was resuspended in 30 mL wash buffer 4 (Table 1) before being sonicated on ice for 30 s on, 30 s off, three times. The suspension was centrifuged at 4 °C at 10 k x g for 15 mins. The washing steps are important to remove impurities from the inclusion bodies and to obtain a high purity of the final recombinant AβM. At this point, the pellet should be white and can be frozen until use or solubilised before chromatography.

### Solubilising and ion exchange chromatography of AβM

The washed inclusion body pellet from 50 mL of culture was placed on ice with a small magnetic stir bar on a magnetic stirrer. 200 μL of 6 M GuHCl was added to the pellet and stirred vigorously for 15 mins to solubilise the AβM containing inclusion bodies. 15 mL of ice cold IEX buffer A (10 mM Tris, 1 mM EDTA, pH 9) was added slowly to the solubilised pellet to dilute the 6M GuHCl and to permit binding of AβM to the ion exchange column. The solubilised AβM was filtered through a 0.22 μm filter before being placed on ice prior to chromatography. If the chromatography system is not kept in a cold room, then all buffers must be kept on ice and the column wrapped in an ice bag to keep the chromatography process as cold as possible to reduce aggregation of AβM. The ion exchange column must be equilibrated prior to the AβM sample being ready for purification to reduce the amount of time of AβM handling. AβM was loaded onto a HiTrap Q HP column (GE, Healthcare) and eluted against a linear gradient of IEX buffer B (10 mM Tris, 1 mM EDTA, 0.75 M NaCl, pH 9) over seven column volumes followed by two column volumes of 100 % buffer B. Purification was performed on an ⍰KTA Pure FPLC and monitored by absorption at 280 nm (GE Healthcare). AβM eluted at ∼30% buffer B and must be immediately placed in aliquots for storage at -80°C. The concentration of AβM was determined by absorption at 280 nm on a NanoVue spectrometer using the extinction coefficient of 1490 M^−1^ cm^−1^ for both AβM variants. AβM42 mass/charge was determined using ESI-MS at the Department of Chemistry, University of Cambridge. To note, for buffer exchange into required buffers for different assays, the fastest method is to use desalting columns and centrifugation. The PD MiniTrap G-10 columns (GE, Healthcare) are suitable for use with small peptides down to 700 Da.

### Expression of recombinant AβM(E22G)-mCherry in HEK293 cells

The plasmid pcDNA5-FRT-TO-Aβ(E22G)-mCherry encodes the arctic mutant of Aβ42 sequence (E22G) and a C-terminus encoded linker sequence GSAGSAAGSGESH followed by the mCherry fluorescent protein sequence. This plasmid was subsequently transfected with the pOG44 plasmid encoding the FLp recombinase into the Flp-In™ T-Rex 293 cell line (#R78007, Thermo Fisher Scientifi. The gene of interest, the coding sequence of AβM(E22G)-mCherry, was integrated into the genome to generate an inducible, stable and single-copy cell line expressing the Arctic mutant (E22G) of Aβ42 fused to mCherry ^15^. Complete media (DMEM (high glucose), 10% FBS and 2 mM L-glutamine) was during cell line construction. Stable transfectants were selected using complete media with the addition of the antibiotic hygromycin B for 6-12 weeks. After selection, single cell clones were collected to generate a homogeneous cell line and the expression level was characterised by flow cytometry in our previous paper ^15^.

To induce AβM(E22G)-mCherry expression, we administered complete media with addition of 1μg/ml tetracycline to the cells. To harvest enough protein for purification, six T75 tissue flasks were seeded and cells induced for protein expression for 7 days. ∼12 million cells were collected and centrifuged at 4000 x g for 15 mins and directly frozen in -80 °C.

### Purification of AβM(E22G)-mCherry

The HEK293 cell pellet was resuspended in 25 mL IEX buffer A (50 mM Tris, pH 8) with protease inhibitors (cOmplete, EDTA-free cocktail, Roche) and sonicated 20 s on, 30 s off, four times using an XL-2020 sonicator (Heat Systems). The suspension was centrifuged at 0.8 K x g for 5 mins at 4°C to remove unbroken cells. The supernatant was removed and centrifuged for a further 15 mins at 21 k x g at 4°C. The supernatant was saved as the soluble fraction and the insoluble fraction was resuspended in IEX buffer A. To determine which fraction contained the most AβM(E22G)-mCherry the fractions were analysed by SDS-PAGE on a 4-12%bis-tris gel and subjected to Western blot analysis to probe for the presence of Aβ and mCherry. The membrane was probed with an anti-Aβ antibody targeted to residues 1-16 (1:1000, #E 610, Biolegend) and a secondary anti-mouse IgG HRP-conjugated antibody (1:1000, #NA931, GE Healthcare). The membrane was dried and reprobed with an anti-mCherry antibody (1:1000, #125096, abcam) and a secondary anti-mouse IgG HRP-conjugated antibody (1:1000, #NA931, GE Healthcare). The soluble fraction was found to contain more AβM(E22G)-mCherry than the insoluble fraction (Supplementary Figure 5.) and was therefore used for further purification. The soluble fraction was filtered through a 0.22 μm membrane before being loaded on a HiScreen Capto Q ImpRes ion exchange column (GE Healthcare). The AβM(E22G)-mCherry was eluted from the column over seven column volumes on a linear gradient against IEX buffer B (50 mM Tris, 1 M NaCl, pH 8) followed by two column volumes of 100% buffer B. Western blot analysis, using the same antibodies as described above, was used to confirm in which eluted fractions the AβM(E22G)-mCherry resided in. Monomeric AβM(E22G)-mCherry has a MW of 32.3 kDa, the Western blots show the presence of both degraded and aggregated AβM(E22G)-mCherry from purification. The AβM(E22G)-mCherry positive fraction was slightly pink in the column, but to note the HEK293 cell medium is also pink and residues of the latter can be present in other eluted fractions, therefore Western blot analysis is required to confirm the presence of AβM(E22G)-mCherry rather than just relying on colour of eluted fractions. The concentration of the eluted fraction containing AβM(E22G)-mCherry was calculated from absorption at 280 nm on a NanoVue spectrometer using the extinction coefficient of 35870 M^−1^ cm^−1^. AβM(E22G)-mCherry mass/charge was determined using ESI-MS at the Department of Chemistry, University of Cambridge.

### SDS-PAGE and Western blot

To determine in which fractions AβM eluted from the ion exchange columns, SDS-PAGE was ran. 20 μL of protein solution was incubated with 4 μL of LDS sample buffer and incubated at 100°C for 5 mins before 10 μL was loaded on a 4-12% bis-tris gel (NuPAGE™, Thermo Fisher Scientific). The gel was either stained with Coomassie blue or transferred to a 0.22 μm polyvinylidene fluoride (PVDF) membrane and probed for Aβ. The membrane was first blocked with 5% BSA in PBS with 0.05% Tween-20 for 30 minutes before incubation with the primary antibody against residues 1-16 of Aβ (1:1000, #E610, biolegend) for one hour. After washing for two mins three times the membrane was incubated with the secondary antibody, anti-mouse IgG linked to HRP (1:1000, #NA931, GE Healthcare) for one hour. The membrane was washed for two minutes five times and incubated with chemiluminescent subtrate (SuperSignal™ WEST pico PLUS, Thermo Fisher Scientific) and imaged using a G:Box (Syngene). The membrane was dried and subsequently reprobed using similar conditions, by blocking with 5% BSA, incubating with the primary antibody against mCherry (1:1000, #[1C51] ab125096, abcam) followed by washing and incubation with the secondary antibody anti-mouse IgG linked to HRP (1:1000, #NA931, GE Healthcare).

### Thioflavin-T (ThT) based kinetic aggregation assays

20 μM freshly made ThT (abcam, Cambridge, UK) was added to 50 μL of 5 μM AβM42 after buffer exchange into 100 mM Tris, 200 mM NaCl pH 7 using PD MiniTrap G-10 columns (GE, Healthcare). All samples were loaded onto nonbinding, clear bottom, 96-well half-area plates (Greiner Bio-One GmbH, Germany). The plates were sealed with a SILVERseal aluminium microplate sealer (Grenier Bio-One GmbH). Fluorescence measurements were taken with a FLUOstar Omega plate reader (BMG LABTECH GmbH, Ortenbery, Germany). The plates were incubated at 37°C with double orbital shaking at 300 rpm for one min before each read every five mins for 600 mins. Excitation was set at 440 nm with 20 flashes and the ThT fluorescence intensity measured at 480 nm emission with a 1300 gain setting. Two ThT assays were run using four fractions of AβM42 from two purification runs with four wells per fraction. Data were normalised to the sample with the maximum fluorescence intensity for each plate.

### Transmission electron microscopy of AβM42 and AβM(E22G)-mCherry aggregates

5 μM of AβM42 was incubated in 100 mM Tris 200 uM NaCl, pH7 for two days with constant rotation at 20 rpm on a rotator (SB2, Stuart Scientific) at 37C. AβM(E22G)-mCherry was used at a concentration of 77 μM. 10 μL of each sample was deposited on a carbon 400 mesh grid for 1 min. The grid was washed twice for 15 s in dH2O before incubating in 2% uranyl acetate for 30 s to negatively stain the sample. The grid was imaged using a Tecnai G2 80-200kv TEM at the Cambridge Advanced Imaging Centre.

### Fluorescence imaging of AβMC40-AF488 and AβM(E22G)-mCherry

A glass coverslip was cleaned with 1 M KOH for 15 mins and washed extensively with dH_2_O and dried. 5 μM of AβMC40-AF488 sample was incubated at 37°C for 2 days with rotation at 20 rpm. The solution was centrifuged at 21 k x g for 5 minutes and 10 μL deposited on the glass and incubated in the dark for 15 mins. 10 μL of AβM(E22G)-mCherry at 77 μM was deposited on the glass and incubated in the dark for 15 mins. Both samples were washed three times to remove unbound protein with dH_2_O and imaged. Images of the samples were collected using a custom-built 3-colour structured illumination microscopy (SIM) setup which we have previously described ^16^. A 60 x/1.2NA water immersion lens (UPLSAPO 60XW, Olympus) and a sCMOS camera (C11440, Hamamatsu) were used. The laser excitation wavelengths used were 488 nm (iBEAM-SMART-488, Toptica) for imaging AβMC40-AF488 and a 561 nm laser for AβM(E22G)-mCherry (OBIS 561, Coherent). The samples were also imaged in the other laser channel and in the 640 nm laser (MLD 640, Cobolt) channel to check for cross talk or non-specific fluorescence contaminants, no/little fluorescence was observed in other channels. The laser intensity used was between 10 and 20 W/cm2 with an exposure time of 150 ms. Although a SIM setup was used the intensity of the signal was too low in the samples to use artefact-free SIM reconstruction, so widefield reconstruction was used and the average intensity from nine SIM images is presented. The same set up was used to image AβM(E22G)-mCherry aggregates in cells, but the signal was strong enough for SIM reconstruction, which was performed with LAG SIM, a custom plugin for Fiji/ImageJ available in the Fiji Updater. LAG SIM provides an interface to the Java 691 functions provided by fairSIM ^17^.

## Author Contributions

A.D.S designed experiments and performed purification, kinetic assays, TEM and widefield microscopy imaging. M.L. constructed the AβM(E22G)-mCherry Flp-In™ T-Rex 293 cell line and performed cell culture, performed SIM imaging, reconstruction of images of AβM(E22G)-mCherry in cells and helped with widefield microscopy imaging. A.D.S and G.S.K.S wrote the manuscript. All authors contributed to the manuscript and gave their final approval.

## Notes

The authors declare no competing financial interests.

## Acknowledgements

We would like to thank Dr Penny Hamyln and Dr Nadezhda Nespovitaya for helpful discussions on chromatography protocols. We also thank Lyn Carter and Filomena Gallo of the Cambridge Advanced Imaging Centre for help with sample loading into the TEM. We thank Dr Dijana Matak-Vinkovic for LC/MS sample analysis at the Department of Chemistry, University of Cambridge. We thank Maria Zachoropoulou for critical analysis of the manuscript. This work was supported by the Wellcome Trust, Alzheimer’s Research UK Grants, the Michael J Fox Foundation and Infinitus China Ltd.

## Supplementary Information

Supplementary contains; **Supplementary Figure 1**. Ion exchange chromatography of AβM42 leads to multimer formation if the AβM42 concentration is high. **Supplementary Table 1**. Concentrations of AβM42 fractions from purification in Figure 2. **Supplementary Figure 2**. The same ion exchange chromatography protocol can be used to purify AβMC40 as AβM42. **Supplementary Figure 3**. AβM42 forms fibrillar structures after incubation. **Supplementary Figure 4**. Mass/charge spectrum of purified recombinant AβM42. **Supplementary Table 2**. Calculation of the concentration and degree of labelling for eluted fractions containing AβMC40-AF488. **Supplementary Figure 5**. Fluorescent fibrillar and oligomeric structures are present after incubation of AβMC40-AF488. **Supplementary Figure 6**. More AβM(E22G)-mCherry is present in the soluble fraction than the insoluble fraction. **Supplementary Figure 7**. Large aggregated structures are present in purified AβE22G-mCherry fractions. **Supplementary Figure 8**. Purified AβM(E22G)-mCherry from HEK293 cells remains fluorescent. **Supplementary Figure 9**. Deconvoluted mass spectrum of AβM(E22G)-mCherry shows many protein species present.

## Supplementary Information

**Supplementary Figure 1.**
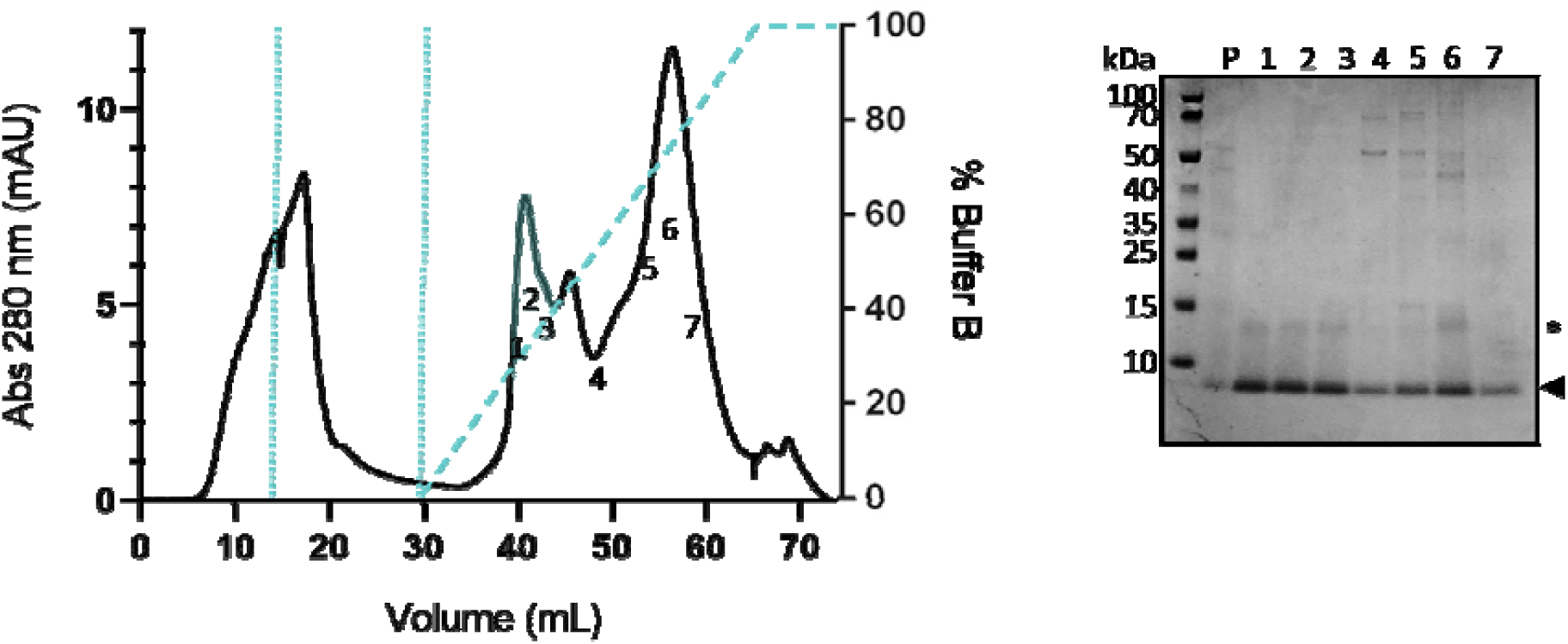
Ion exchange chromatography of AβM42 leads to multimer formation if the AβM42 concentration is high. (a.) Solublised AβM42 was loaded onto a HiTrap Q HP ion exchange column (up to the first dotted line in a.) and the unbound protein washed from the column (up to the second dotten line in a.). AβM42 was eluted along a linear gradient of buffer B containing 0.75 M NaCl (dashed line showing gradient in a.). (b.) To determine where AβM42 eluted from the column the fractions collected were analysed using SDS-PAGE on a 4-12% bis-tris gel and stained with Coomassie blue. The Aβ sample prior to IEX (P) contained some contaminants. Protein bands correlating to ∼ 4.5 kDa (shown by the arrow next to b.) show monomeric AβM42 in all fractions. The most pure fractions 1-3 are highlighted in blue in the chromatograph (a.), however there were some multimeric structures present (indicated by the star) suggesting that this sample prep or purification run was not performed rapidly enough to obtain solely monomeric protein. AβM42 eluted at ∼ 30% buffer B.

**Supplementary Table 1.** Concentrations of AβM42 fractions from purification in Figure 2.

| **Fraction** | **Concentration (μM)** |
| --- | --- |
| 2 | 8 |
| 3 | 11.4 |
| 4 | 12.75 |
| 5 | 12.75 |
| 6 | 12.75 |
| 7 | 12 |

**Supplementary Figure 2.**
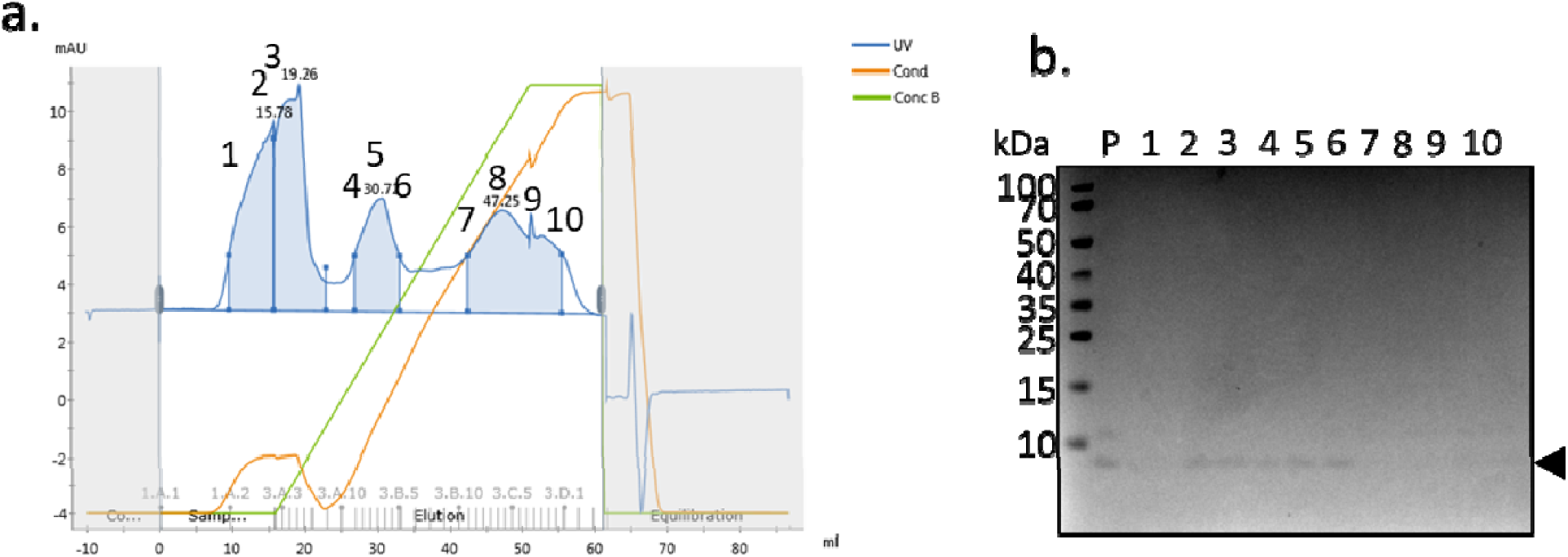
The same ion exchange chromatography protocol can be used to purify AβMC40 and AβM42. (a.) Solublised AβMC40 was loaded onto a HiTrap Q HP ion exchange column and was eluted along a linear gradient of buffer B containing 0.75 M NaCl (green line). The increase in salt concentration was monitored by conductivity (orange line) and absorption at 280 nm was used to detect the protein eluting from the column (blue line) (b.) To determine where AβMC40 eluted from the column the fractions collected were analysed using SDS-PAGE on a 4-12% bis-tris gel and stained with Coomassie blue. The Aβ sample prior to IEX (P) contained some contaminants. Protein bands correlating to ∼ 4.5 kDa (shown by the arrow next to b.) show monomeric AβMC40 in all fractions 3-7.

**Supplementary Figure 3.**
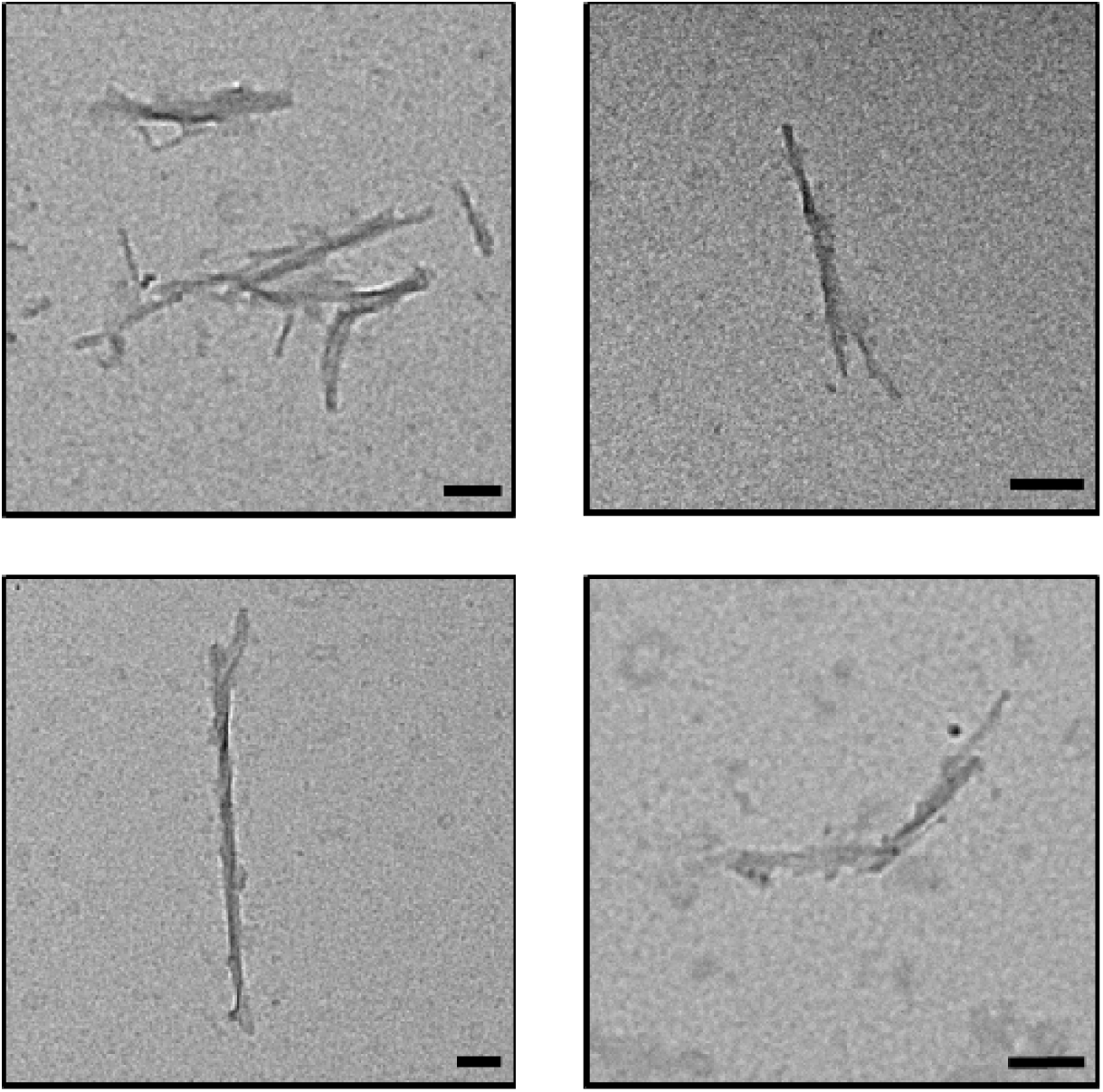
AβM42 forms fibrillar structures after incubation. Fibrils formed during incubation of 5 μM of AβM42 at 37°C with constant rotation at 20 rpm for two days. 10 μL of sample was incubated on a 400 mesh carbon coated copper grid and negatively stained with 2% uranyl acetate. Scale bar = 100 nm.

**Supplementary Figure 4.**
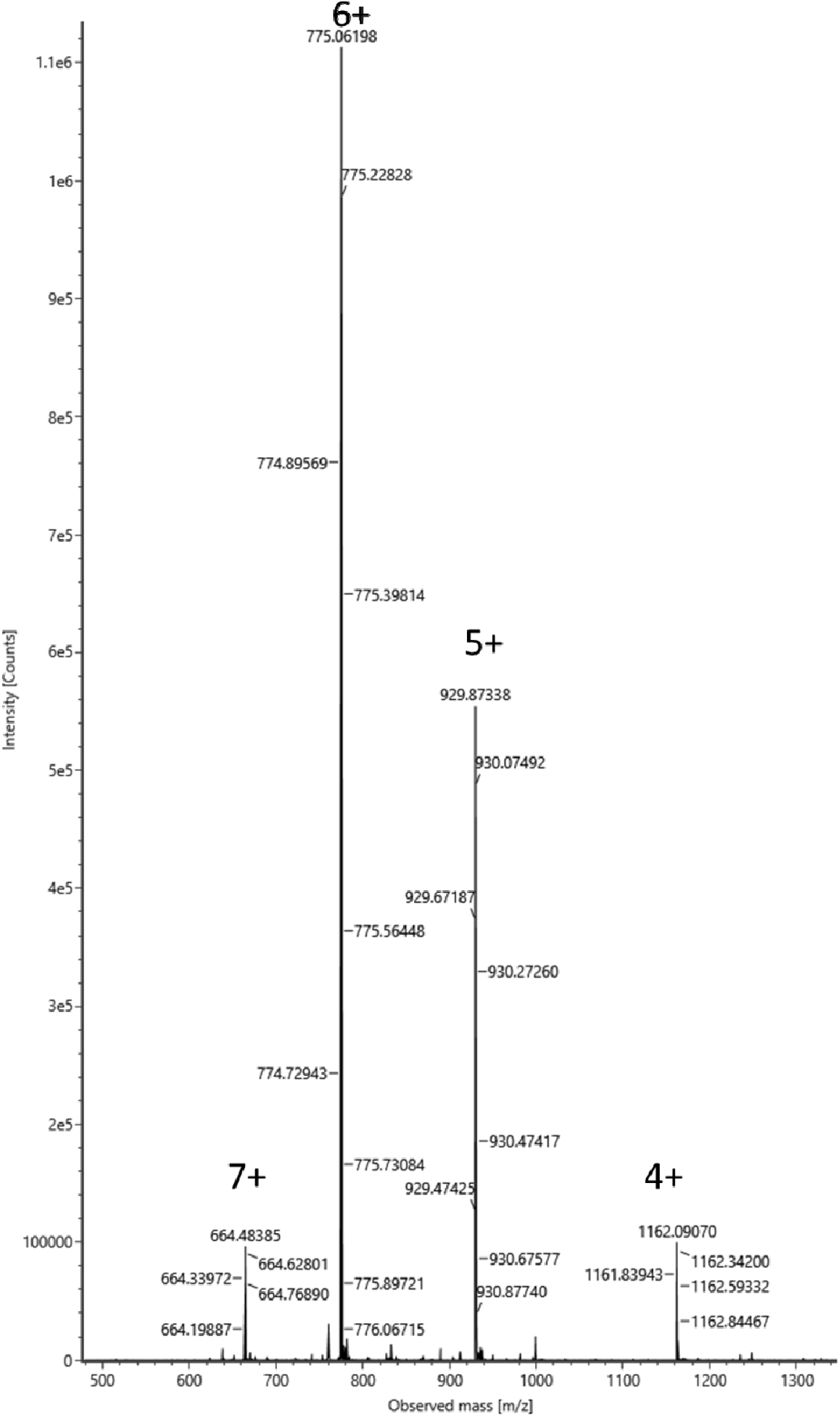
Mass/charge spectrum of purified recombinant AβM42. MS of AβM42 displaying [M+7H]^7+^ at 664.48 m/z, [M+6H]^6+^ at 775.06 m/z, [M+5H]^5+^ at 929.87 m/z and [M+4H]^4+^ at 1162.09 m/z.

**Supplementary Table 2.**
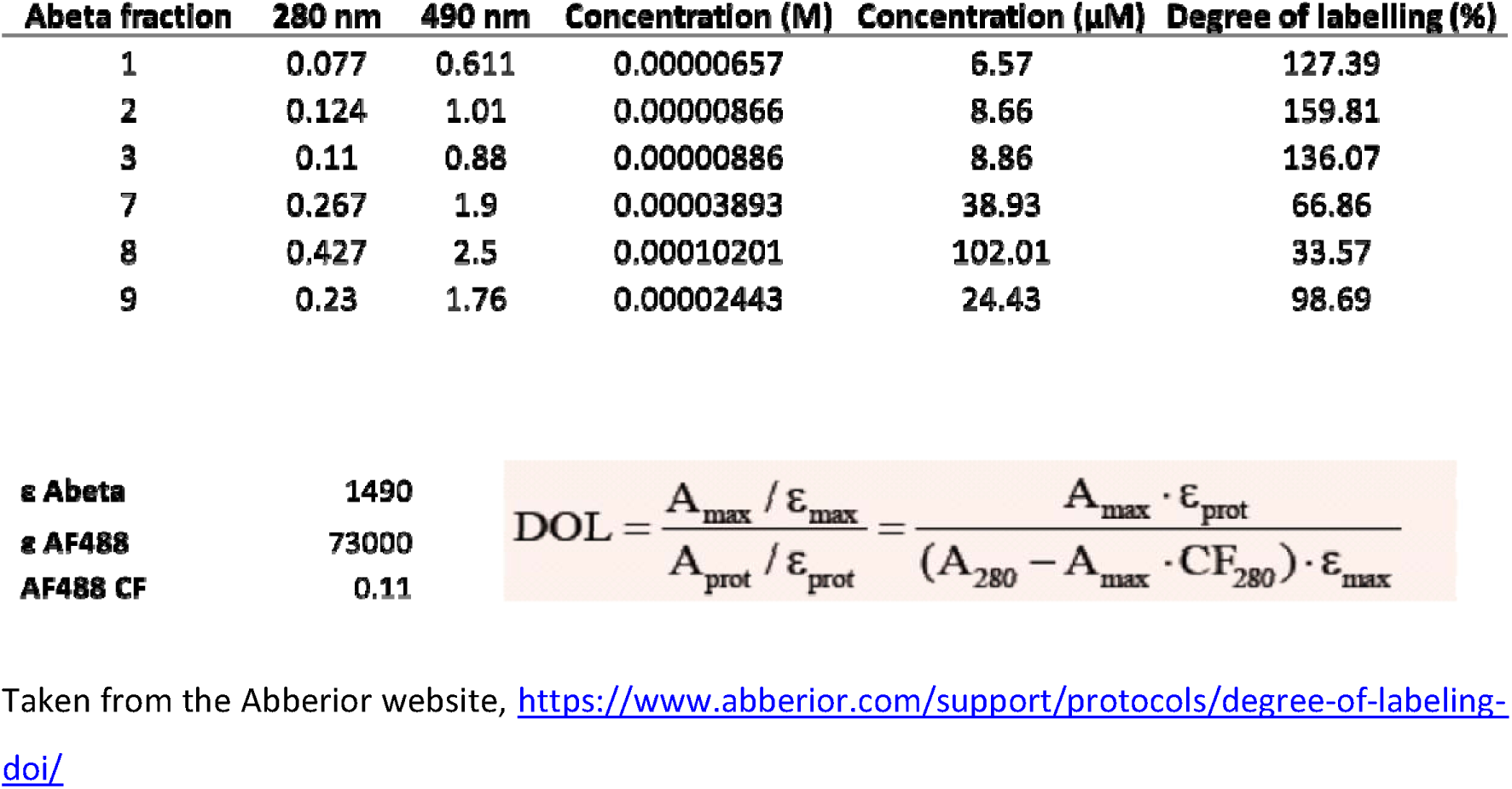
Calculation of the concentration and degree of labelling for eluted fractions containing AβMC40-AF488.

| Abeta fraction | 280 nm | 490 nm | Concentration (M) | Concentration (μM) | Degree of labelling (%) |
| --- | --- | --- | --- | --- | --- |
| 1 | 0.077 | 0.611 | 0.00000657 | 6.57 | 127.39 |
| 2 | 0.124 | 1.01 | 0.00000866 | 8.66 | 159.81 |
| 3 | 0.11 | 0.88 | 0.00000886 | 8.86 | 136.07 |
| 7 | 0.267 | 1.9 | 0.00003893 | 38.93 | 66.86 |
| 8 | 0.427 | 2.5 | 0.00010201 | 102.01 | 33.57 |
| 9 | 0.23 | 1.76 | 0.00002443 | 24.43 | 98.69 |

|  |  |  |
| --- | --- | --- |
| **ε Abeta** | 1490 | $\text{DOL} = \frac{A_{\text{max}} / \epsilon_{\text{max}}}{A_{\text{prot}} / \epsilon_{\text{prot}}} = \frac{A_{\text{max}} \cdot \epsilon_{\text{prot}}}{(A_{280} - A_{\text{max}} \cdot \text{CF}_{280}) \cdot \epsilon_{\text{max}}}$ |
| **ε AF488** | 73000 |  |
| **AF488 CF** | 0.11 |  |

Taken from the Abberior website, https://www.abberior.com/support/protocols/degree-of-labeling-doi/

**Supplementary Figure 5.**
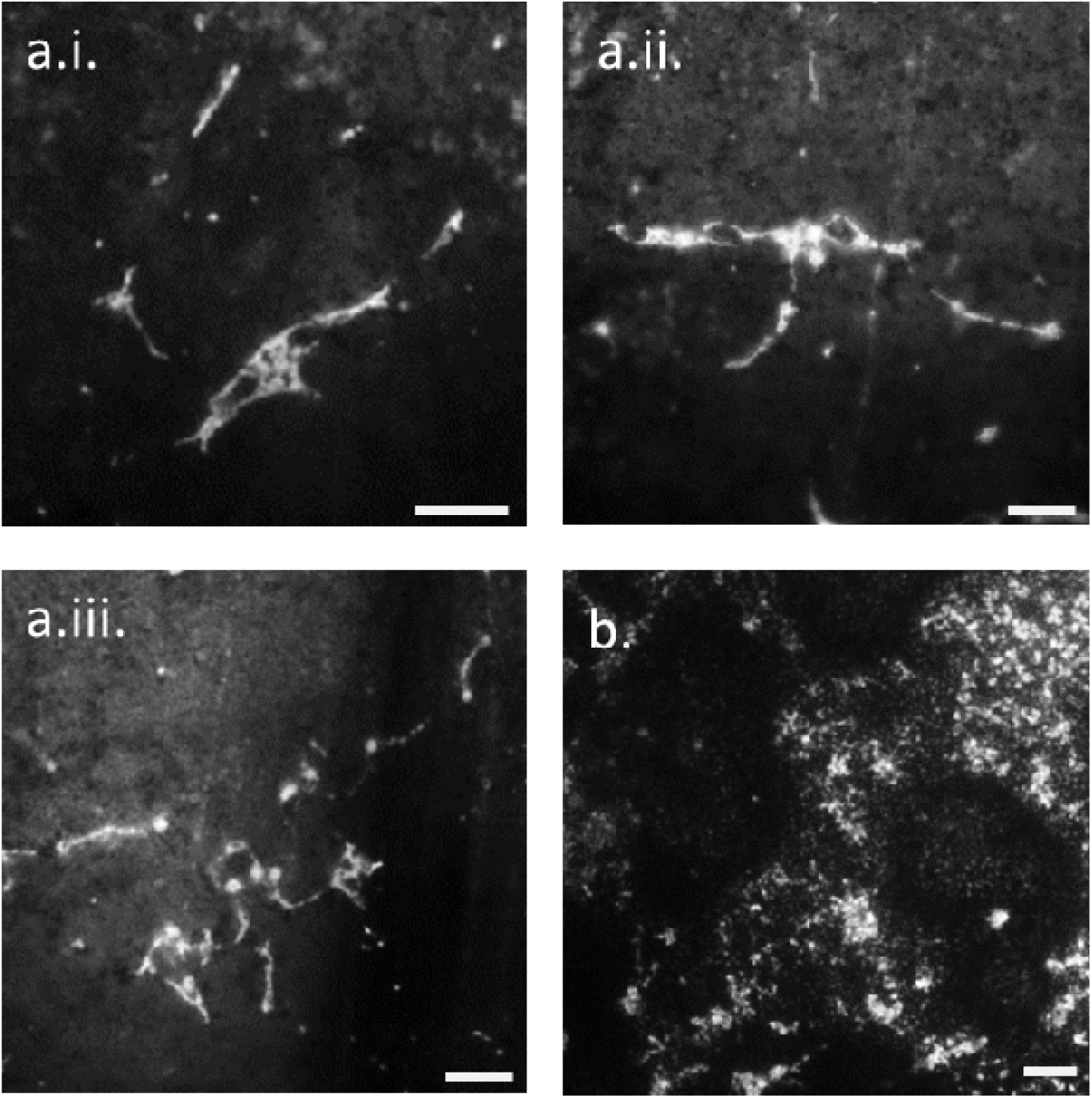
Fluorescent fibrillar and oligomeric structures are present after incubation of AβMC40-AF488. 5 μM of 100% labelled AβMC40-AF488 was incubated for two days at 37°C at 20 rpm. The sample was deposited on a glass coverslip and excited with a 488 nm laser and (a.i-iii.) fluorescent fibrils and (b) oligomeric structures were observed using a widefield microscope. Scale bar 5 μm.

**Supplementary Figure 6.**
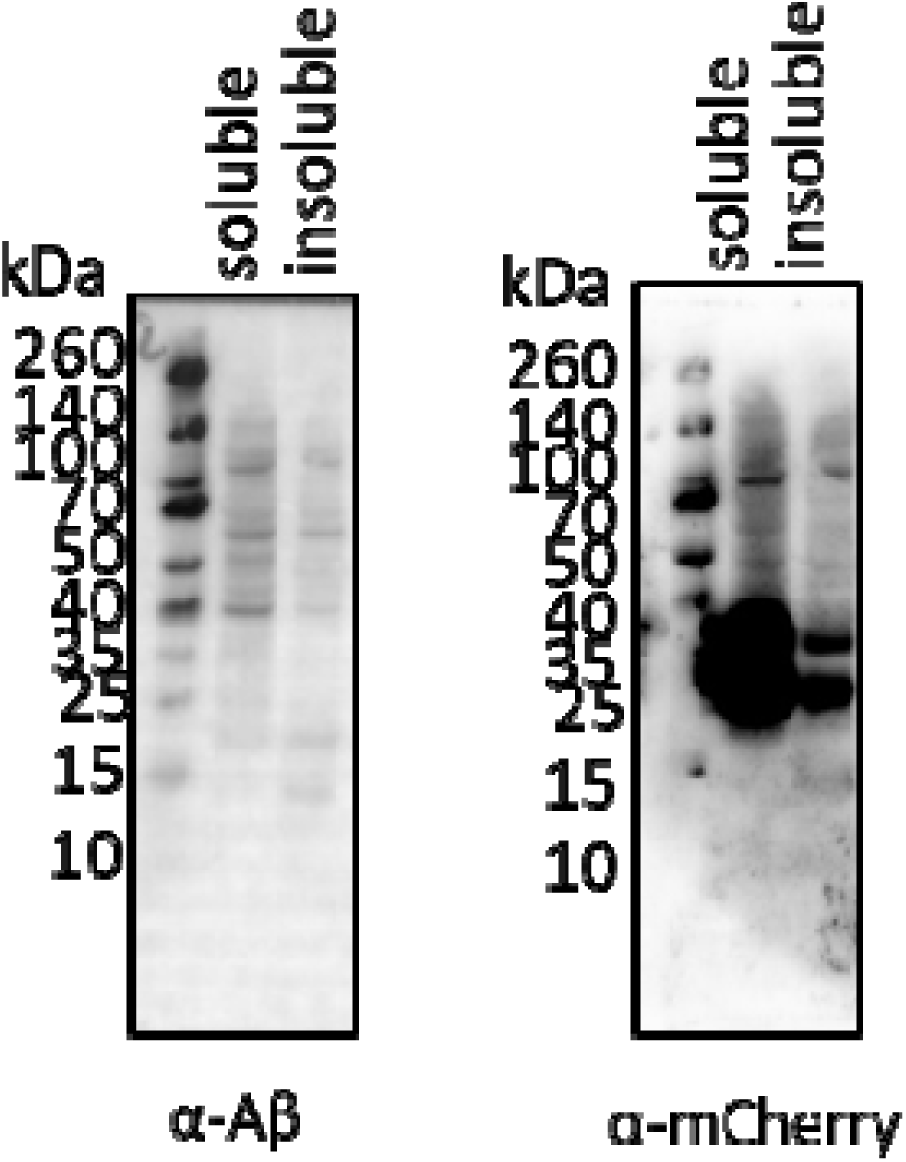
More AβM(E22G)-mCherry is present in the soluble fraction than the insoluble fraction. AβM(E22G)-mCherry expressed in HEK293 cells was lysed from the cell by sonication and centrifuged to separate the soluble and insoluble fractions. Analysis of the two fractions by Western blot using an anti-Aβ antibody (E610) (α-Aβ) and reprobing the same membrane with an anti-mCherry antibody (α –mCherry) shows more AβM(E22G)-mCherry in the soluble fraction.

**Supplementary Figure 7.**
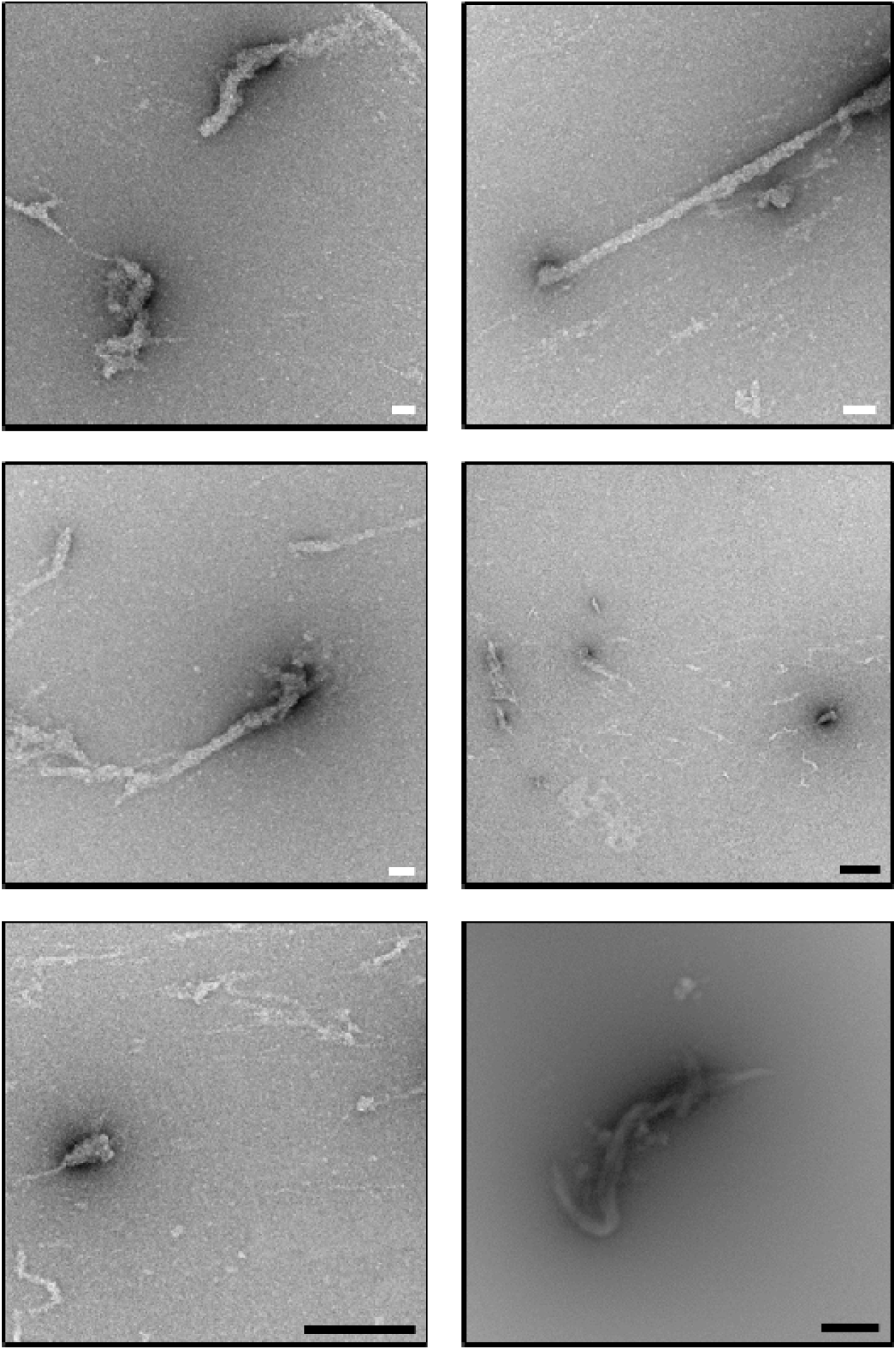
Large aggregated structures are present in purified AβE22G-mCherry fractions. 10 μL of AβE22G-mCherry sample was incubated on a 400 mesh carbon coated copper grid and negatively stained with 2% uranyl acetate before imaging with TEM. White scale bar = 100 nm, black scale bar = 500 nm.

**Supplementary Figure 8.**
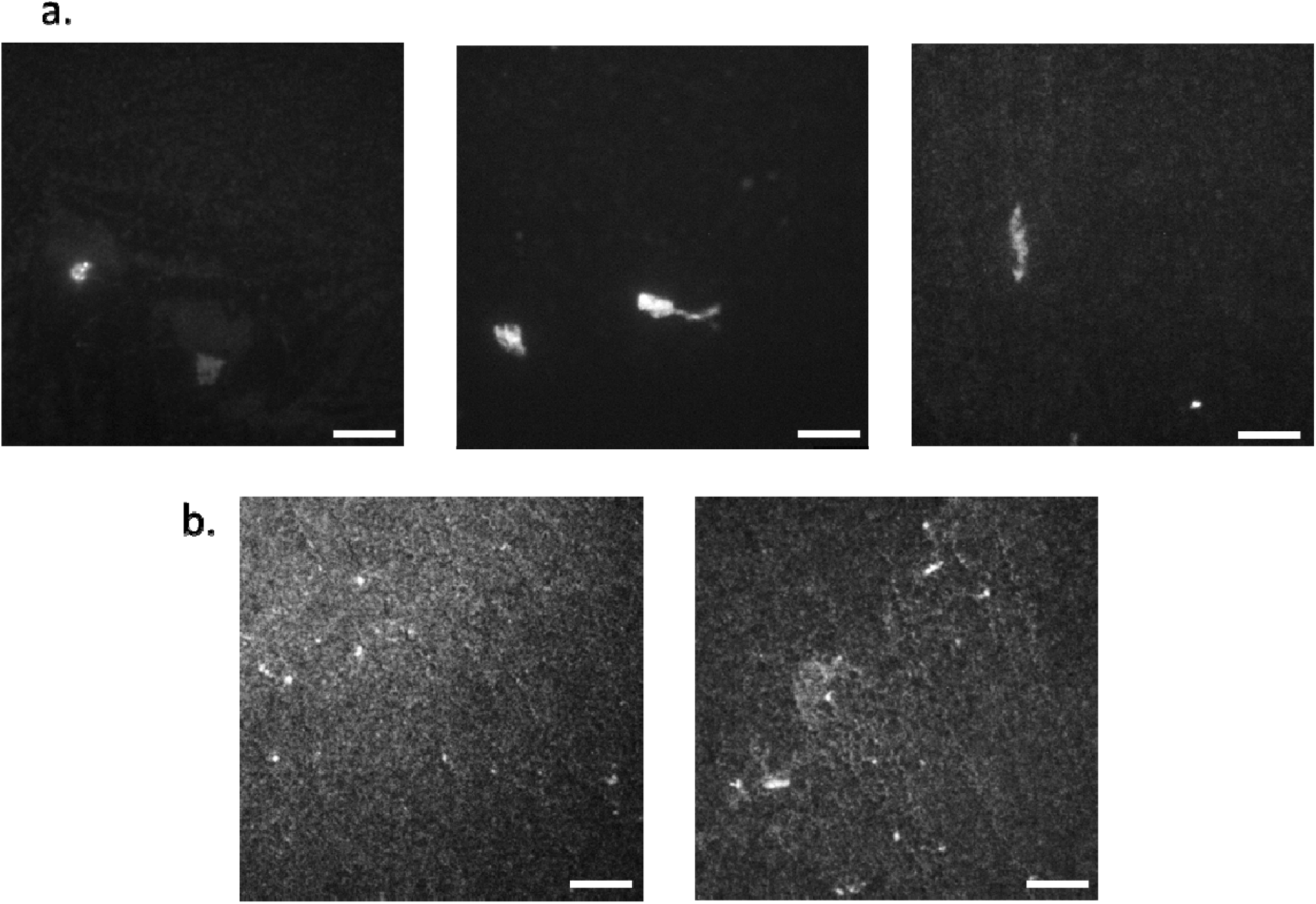
Purified AβE22G-mCherry from HEK293 cells remains fluorescent. AβE22G-mCherry isolated from HEK293 cells and purified by ion exchange chromatography was incubated on a glass coverslip and imaged with widefield microscopy. A 561 nm laser was used to excite the sample and (a.) large aggregated structures and (b.) small aggregated structures displayed fluorescence. Scale bar = 8 μm.

**Supplementary Figure 9.**
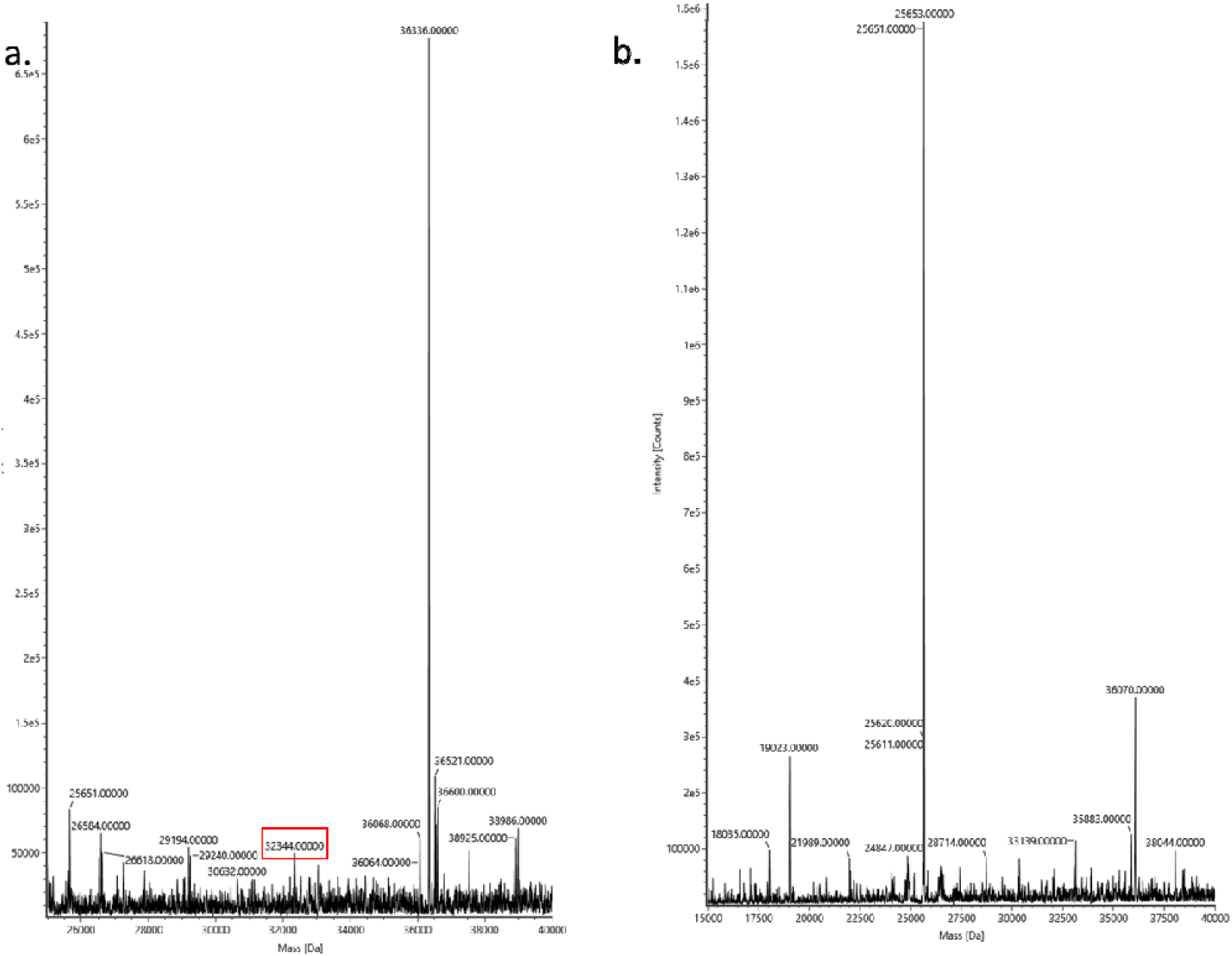
Deconvoluted mass spectrum of AβM(E22G)-mCherry shows many protein species present. The AβM(E22G)-mCherry sample displayed multiple peaks on the chromatograph prior to MS, two fractions were electrosprayed for MS analysis (a. (eluting at 4. 5 mins) and b (eluting at 4.9 mins)). The expected MW of AβM(E22G)-mCherry is ∼32.3 kDa (highlighted in the red box in a.), however this is not the most abundant species isolated from HEK293 cells. A larger species of ∼36.3 kDa (a.) and a degraded product of ∼25.6 kDa (b.) are instead the dominant protein species by MS. Also present are many other species of differing MW.

